# Uncovering complementary sets of variants for predicting quantitative phenotypes

**DOI:** 10.1101/2020.12.11.419952

**Authors:** Serhan Yılmaz, Mohamad Fakhouri, Mehmet Koyutürk, A. Ercüment Çiçek, Öznur Taştan

**Affiliations:** Department of Computer and Data Sciences, Case Western Reserve University; Department of Computer Engineering, Bilkent University; Center for Proteomics and Bioinformatics, Case Western Reserve University; Department of Computational Biology, Carnegie Mellon University; Faculty of Engineering and Natural Sciences, Sabanci University

## Abstract

**Motivation:** Genome-wide association studies show that variants in individual genomic loci alone are not sufficient to explain the heritability of complex, quantitative phenotypes. Many computational methods have been developed to address this issue by considering subsets of loci that can collectively predict the phenotype. This problem can be considered a challenging instance of feature selection in which the number of dimensions (loci that are screened) is much larger than the number of samples. While currently available methods can achieve decent phenotype prediction performance, they either do not scale to large datasets or have parameters that require extensive tuning.

**Results:** We propose a fast and simple algorithm, Macarons, to select a small, complementary subset of variants by avoiding redundant pairs that are in linkage disequilibrium. Our method features two interpretable parameters that control the time/performance trade-off without requiring parameter tuning. In our computational experiments, we show that Macarons consistently achieves similar or better prediction performance than state-of-the-art selection methods while having a simpler premise and being at least 2 orders of magnitude faster. Overall, Macarons can seamlessly scale to the human genome with ~10^7^ variants in a matter of minutes while taking the dependencies between the variants into account.

**Conclusion:** Macarons can offer a reasonable trade-off between phenotype predictivity, runtime and the complementarity of the selected subsets. The framework we present can be generalized to other high-dimensional feature selection problems within and beyond biomedical applications.

**Availability:** Macarons is implemented in Matlab and the source code is available at: https://github.com/serhan-yilmaz/macarons

## 1 Introduction

Genome-Wide Association Studies (GWAS) attempt to find a relation between the genetic variations and a phenotype. Many single nucleotide polymorphisms (SNPs) have been found to be associated with various diseases or disorders including type II diabetes, obesity and schizophrenia as well as other quantitative traits like height individually [12, 32]. However, individual SNPs fail to explain complex phenotypes, in which multiple SNPs contribute collectively [20]. Thus, as an alternative and more powerful approach, many studies aim at finding a good subset of SNPs that are associated with the phenotype of interest as a group [33, 5, 26, 34]. This study is mainly focused on the problem of finding a subset of SNPs that are *collectively* predictive of the phenotype of interest. For the sake of brevity, we will simply refer to it as the SNP subset selection problem throughout this paper.

### Approaches investigating combinations of SNPs

Finding combinations of SNPs that are predictive of a phenotype is computationally challenging due to the large number of possible combinations that need to be considered. There are methods that focus on high-order interactions using exhaustive search [25, 19] or greedy algorithms [10, 37] on a small, limited pool of SNPs that is usually not more than a few hundreds [29, 11]. While such pools of ”promising SNP candidates” are typically obtained using *a priori* information sources by limiting the analysis to SNPs residing in the coding regions of the genome, it is also possible to conduct a filtering based on automated searches [31, 9].

Indeed, for combinatorial studies investigating the pair-wise interactions (or as more commonly known as *epistasis*) between the variants in full genome, recent studies show the importance of limiting the search space through prioritization of the tests [27, 6], both to alleviate the computational intensity of the task, as well as to improve the overall statistical power. Specifically, Caylak *et al*. [4] demonstrates the utility of an initial filtering of SNPs based on an automated SNP selection algorithm [36] as a powerful approach that can improve the statistical power considerably.

### Approaches for SNP selection problem in quantitative phenotypes

The SNP selection problem in quantitative phenotypes essentially corresponds to a feature subset selection problem for multivariate regression [23]. However, due to the high dimensional nature of typical GWAS data (millions of variants), established methods for feature selection such as linear regression with *l*_1_ (lasso) regularization [30, 13], spectral-relaxation based approaches [38, 8], graph-constrained feature selection methods like GraphLasso and GroupLasso [22, 15], as well as various other methods with sparsity constraints known in the bioinformatics community in the context of other problems (e.g., for selecting gene sets) [17, 16, 18], are computationally too expensive for this task. Thus, a common strategy is to apply a simple threshold-based filtering (e.g., a p-value cutoff) based on individual phenotype associations [31], for example, using a statistical test like sequence kernel association test (SKAT) [35]. The downside of this approach is that threshold-based filtering considers each variant independently and does not take into account of the dependencies or interactions between them.

To achieve a scalable solution for all known variants in the genome while considering the dependencies between them, alternative SNP selection algorithms have been proposed [3, 36]. Such algorithms simplify the problem by focusing on a linear combination of individual phenotype associations of SNPs while using some *a priori* information encoded in the form of a biological network to improve the overall predictivity of the selected subset. In particular, SConES [3] uses a minimum-cut solution under sparsity and connectivity constraints on a SNP-SNP network. More recently, SPADIS [36] selects a diverse set of SNPs using the SNP-SNP network.

### The drawbacks of existing methods

Linkage disequilibrium (LD) which refers to the non-random association of variants, is a common phenomenon for close variants on the same chromosome [1]. While the connectivity constraint of SConES helps to improve the quality of the selected set, it implicitly promotes the selection of SNPs that are in LD impairing the prediction performance. On the other hand, SPADIS seeks to increase the diversity of SNPs by penalizing the selection of close SNPs on the input network. While this diversity helps to avoid redundant SNPs in LD and improves the phenotype predictions, the drawback of SPADIS is that it requires two parameters without any interpretable meanings or default values, that need to be tuned through an external procedure such as cross-validation. The need for such external procedures not only makes the method hard to apply from a user viewpoint, but also considerably exacerbates the run time and reduces the robustness of the selections when there are time and resource constraints.

### Macarons: a fast and simple algorithm to select complementary SNPs

To overcome these limitations, we determined three main objectives a SNP selection algorithm should satisfy: (i) Have good prediction performance for quantitative phenotypes (at least as predictive as available methods), (ii) Fast enough to consider all variants in the genome, and (iii) Easy to use without requiring external parameter tuning procedures like cross-validation. Thus, we propose a new algorithm named Macarons that take into account the correlations between SNPs to avoid the selection of redundant pairs of SNPs in linkage disequilibrium. Overall, Macarons features two simple, interpretable parameters to control the time/performance trade-off: the number of SNPs to be selected (k), and maximum intra-chromosomal distance (*D*, in base pairs) to reduce the search space for redundant SNPs. Note that, since the parameters have interpretable meanings, they can be determined in advance (without requiring an external procedure for parameter tuning) with the available computational resources and the goals of further studies in mind.

## 2 Methods

### 2.1 Background

#### 2.1.1 Problem Definition

We are given as input a ground set of SNPs *V* of cardinality *n*, genotype matrix **X** ∈ {0,1,2}^*m×n*^ decoding the number of alternate alleles for *m* samples and n SNPs, and a phenotype vector 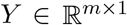 containing quantitative values for m samples. The number of SNPs n is much larger than the number of samples *n* ≫ *m*. Thus, we would like a obtain a small subset of SNPs *S* = {*s*_1_, *s*_2_,…, *s_k_*} ⊆ *V* of size *k* that maximizes the prediction performance of the given phenotype vector *Y* based on a regression model 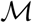. In this study, we consider a linear model (i.e., without interaction terms modeling epistasis), where each selected SNP *s_i_* ∈ *S* has an additive effect on the phenotype:

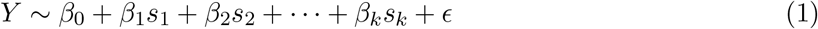

where *β_i_* is the regression coefficients to be learned from data and *ϵ_i_* is an error term that is normally distributed with zero mean. Based on this model, the collective effect of the SNP set S can be characterized by the squared multiple correlation coefficient *R*^2^(*Y, S*) which has the interpretation of the variance explained in *Y* by *S*. Thus, the overall SNP selection problem can be defined as a SNP subset search problem that maximize the following function:

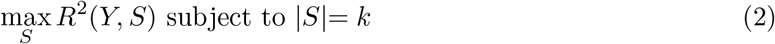

**Algorithm 1.**
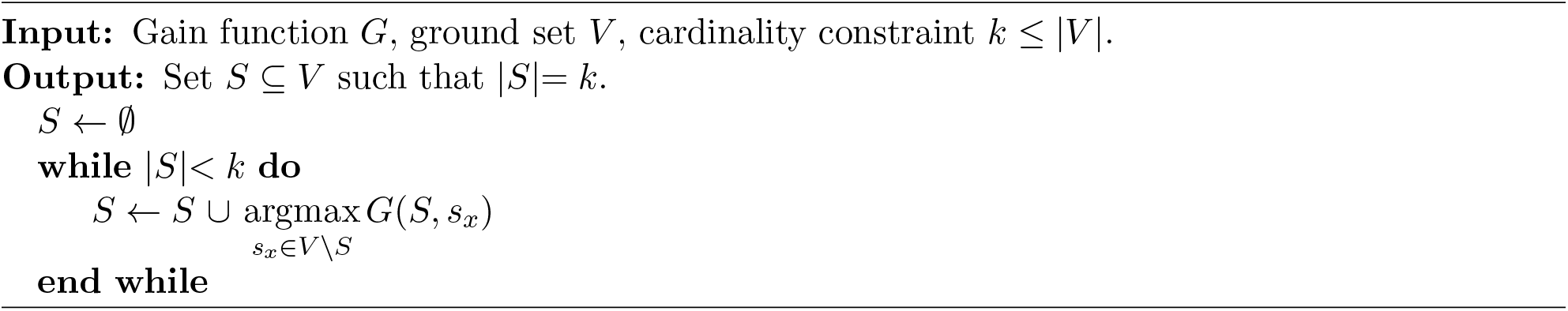
Greedy Subset Selection Algorithm

#### 2.1.2 Forward Step-wise Regression

Generally, solving the regression problem given in Equation 2 is NP-hard [24]. However, due to near submodularity of *R*^2^, greedy formulations that iteratively grow a set based on a local gain function *G* (as in Algorithm 1), produce near-optimal results, proving a good approximation for maximizing *R*^2^ under a cardinality constraint [7, 8]. Among such algorithms, a notable one that is commonly used is the forward step-wise regression that maximizes semi-partial squared correlation as its gain function:

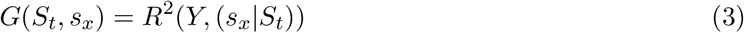

where *S_t_* is the subset of selected features at the *t* iteration of the algorithm, *s_x_* is a candidate feature being considered, and *R*^2^(*Y*, (*s_x_*|*S_t_*)) is the semi-partial correlation coefficient between *Y* and *s_x_*|*S_t_*, when *s_x_* is regressed and residualized with every variable in *S_t_*.

The main issue with using this approach for the SNP selection problem is that it requires estimating and inverting the covariance matrix. This requirement not only makes the algorithm computationally intensive with *O*(*n*^3^) runtime complexity, but also leads to the selection of SNP sets that is likely to overfit to the given training data (this is due to the high dimensionality nature of the problem where *n ≫ m*).

### 2.2 Macarons

Here, we follow an approach similar to the forward step-wise regression where we iteratively grow the selected SNP set based on their estimated contribution *G* for phenotype prediction as measured by the semi-partial correlation. However, to scale to all SNPs in a typical GWAS study as well as to improve the robustness of the algorithm, we apply some simplifying assumptions that reduce the computational complexity and error in estimation. First, we start by expressing the semi-partial correlation *R*^2^(*Y*, (*s_x_*|*S_t_*)) in an alternate form:

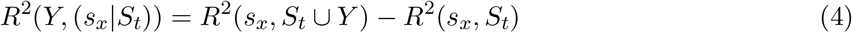

where *R*^2^(*s_x_, S_t_* ∪ *Y*) and *R*^2^(*s_x_, S_t_*) are multiple correlation terms corresponding to linear models predicting *s_x_* using the SNP set *S_t_* with and without the phenotype variable *Y* respectively. Here, we can further decompose *R*^2^(*s_x_, S_t_* ∪ *Y*)into two parts:

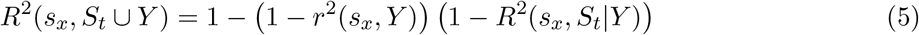

where *r*^2^(*s_x_,Y*) is the squared Pearson’s correlation coefficient indicating the individual predictivity of *s_x_* on *Y*, and *R*^2^(*s_x_, S_t_*|*Y*) is the partial correlation between *s_x_* and *S_t_* given *Y*. Here, we assume that the portion of variance that overlap between *s_x_* and *S_t_* does not depend on their overlap with *Y*, which can be expressed as follows:

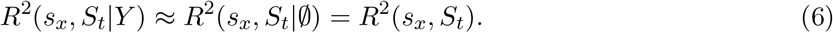

With this assumption, the gain function *R*^2^(*Y*, (*s_x_*|*S_t_*)) can be simplified as follows:

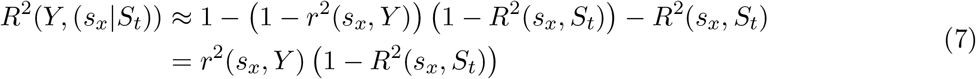

Here, since *r*^2^(*s_x_,Y*) quantifies the individual predictivity of the candidate SNP *s_x_* on phenotype *Y*, which we can also replace with other phenotype association scores (denoted *c_x_* for SNP *s_x_*) such as sequence kernel association test (SKAT). Thus, a more general gain function can be defined as follows:

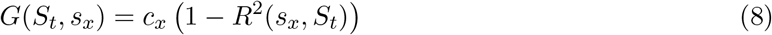

Overall, the multiple correlation *R*^2^(*s_x_, S_t_*) measures the collective redundancy between *s_x_* and *S_t_*, and here used as a penalty function to facilitate the selection of complementary SNPs for the the phenotype prediction.

#### 2.2.1 Estimating the penalization function

The main challenge in estimating the multiple correlation is that it requires the computation of high order interaction terms among the selected SNPs. This makes its estimation for a given data sample both computationally intensive (with *O*(*mt*^2^ +*t*^3^) runtime complexity), and noisy. To help overcome these issues, we first express the multiple correlation as multiplication of several terms involving partial correlations:

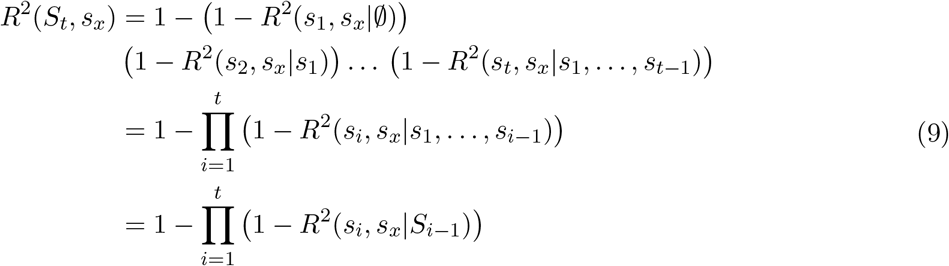

where *R*^2^(*s_i_, s_j_*|*S′*) denotes the squared partial correlation between SNPs *s_i_* and *s_j_* given the SNPs within the set *S′*. Note that, these partial correlation calculations also require computing high-order interactions; thus, do not simplify the computation of the multiple correlation by themselves. For this purpose, we make the following simplifying assumption:

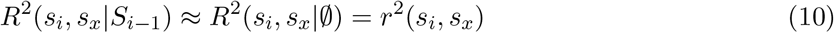

where *r*^2^(*s_i_, s_x_*) is the squared zero-order correlation coefficient (i.e., ordinary Pearson’s correlation) between SNPs *s_i_* and *s_x_*. Thus, with this assumption, the estimation of the multiple correlation simplifies to:

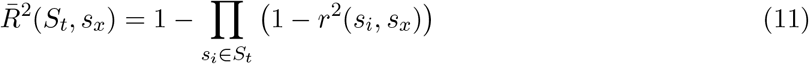

This assumption helps with the overfitting problem in the estimation of multiple correlation since it reduces the number of parameters needed to be estimated from the data and reduces the required computation time drastically (from *O*(*mt*^2^ + *t*^3^) to *O*(*mt*)).

In the remaining sections of this manuscript, we will refer to the estimation of the squared multiple correlation 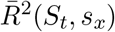 simply as the penalization function, and we will refer to the zero-order correlation *r*^2^(*s_i_, s_x_*) as the redundancy function.

Similar to the squared multiple correlation *R*^2^(*S_t_, s_x_*), this simplified penalization function has several useful properties such as being bounded in [0,1] region, being monotonic, and applying diminishing returns principle (where the increase in penalization decreases proportionally on subsequent iterations as the selected set grows). We explain these properties in more detail in Supplementary Text 1.

#### 2.2.2 Limiting the search space through intra-chromosomal distance

One particular issue for directly using the penalization function given in Equation 11 together with the gain function and algorithm in Equation 8 and Algorithm 1 is that the overall runtime can still be slow for large *k* (number of SNPs selected) with algorithmic complexity of *O*(*nmk*) due to the requirement of computing *O*(*nk*) correlation coefficients. For this reason, we make an additional simplifying assumption to limit the search space to intra-chromosomal SNP pairs within a specified distance. Specifically, we assume the following:

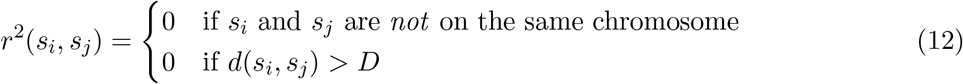

where *d*(*s_i_, s_j_*) is defined as the intra-chromosomal distance between SNPs *s_i_* and *S_j_* (i.e., the distance on the genome) and *D* is an adjustable parameter (unit in base pairs) to control the time/performance trade-off of the algorithm by limiting the search space for the redundancy estimations. Note that, we consider the *d*(*s_i_, S_j_*) to be infinite for SNP pairs that are on different chromosomes.

#### 2.2.3 Formulation of Macarons algorithm

Overall, with the three assumptions given in Equation 6, Equation 10, and Equation 12, the penalty function becomes as follows:

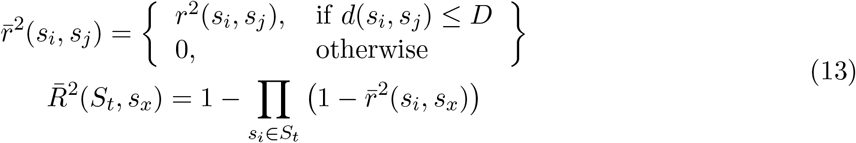

Thus, the gain function becomes:

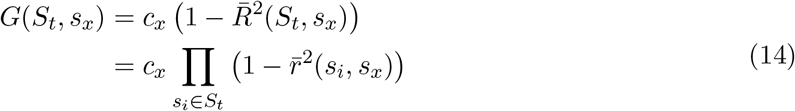

The Macarons algorithm that encodes this gain function for step-wise SNP selection is given in Algorithm 2. Overall, it has *O*(*nk* + λ_*D*_*mk*) run time complexity where the first term is for maximizing the gain function, and the second term is for computing the gain function, which require the measurement of correlations from data. Here, λ_*D*_ is a variable between [1, *n*] dependent on the *D* parameter. It represents the average number of SNP pairs that require the computation of correlation for a given *D* threshold. Thus, the overall complexity for small *D* is *O*(*nk*) when the first term dominates and *O*(*nmk*) for large *D* as the computation of Pearson correlations becomes the bottleneck.

**Algorithm 2.**
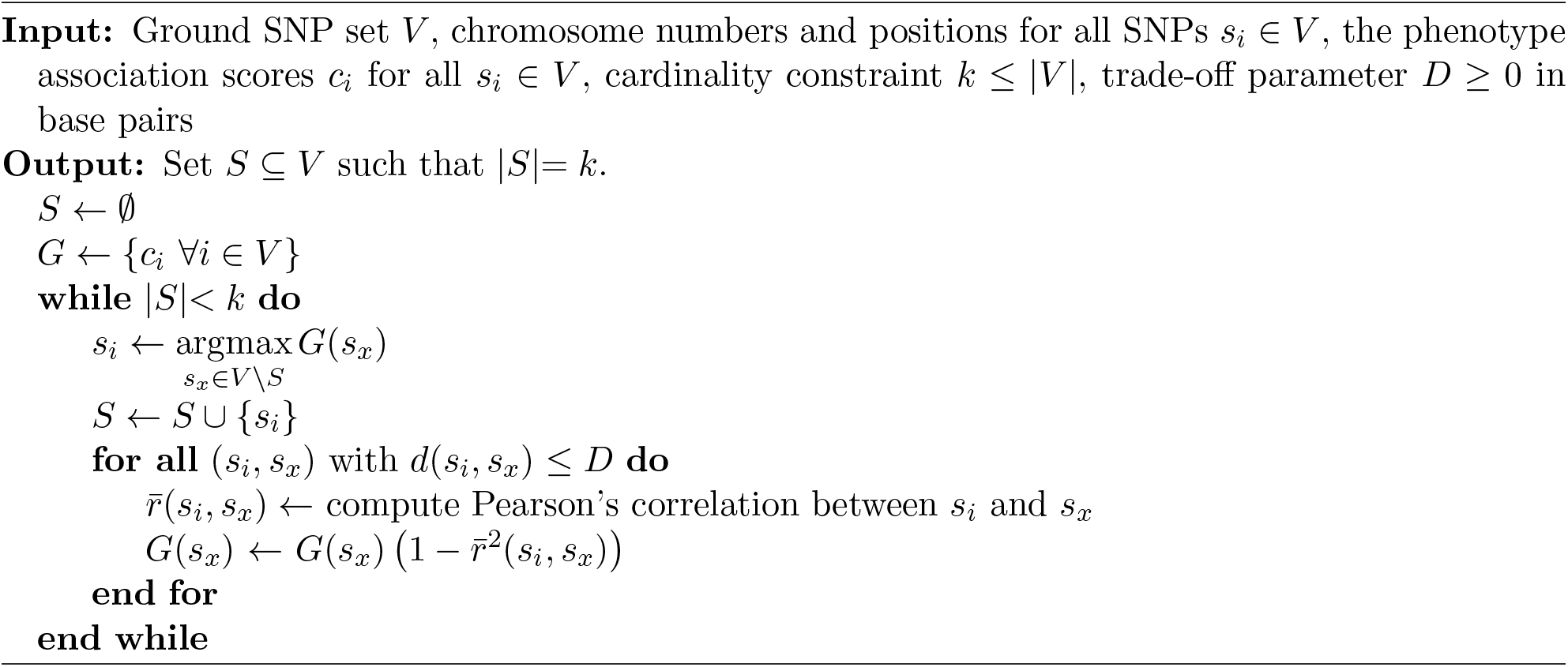
Macarons Algorithm

#### 2.2.4 Optimizing Macarons algorithm for runtime

The gain function given in Equation 14 is monotonically non-increasing with respect to the growing set of selected SNPs (i.e., at each iteration, the gain of a SNP either stays the same or decreases). Moreover, we know that the selected SNP set will approximately grow according to their individual association scores (this is particularly true for low *k* and *D* parameters since there would be less deviation from individual scores). Here, we leverage these properties to further optimize the runtime of the algorithm. For this purpose, we first sort all SNPs according to their phenotype association scores *c_x_* (such that *c_i_* ≥ *C_j_* if *i < j*). Then, we limit the search space of the algorithm to an *active region* consisting of Native most promising SNPs with highest individual scores (having an initial size of *N*_active_ = *ψ*). When the current search space is insufficient (this can be detected by comparing the gain function with the individual scores), we grow the active region by a factor of *γ* > 1. Specifically, when the maximum value of gain function is greater than or equal to the minimum individual score in the active region (i.e., when max_*x≤N*_active__(*G*(*S_t_, s_x_*)) ≥ min_*x≤N*_active__(*c_x_*)), we know that active region is sufficient (since we know min_*x≤N*_active__(*c_x_*) > *C_j_* ≥ *G*(*S_t_*, *S_j_*) ∀{*j* > *N*_active_}). Otherwise, the action region might be insufficient, thus, we grow the active region to include the most promising ⌈*γN*_active_⌉ SNPs and repeat this process as necessary. The optimized Macarons algorithm that implements this idea is given in Algorithm 3. Note that, the output of this algorithm is always equal to the output Algorithm 2 regardless of the parameter values (i.e., the parameters *ψ* and *γ* does not change the output, only affects the runtime). In our experiments, we use *ψ* = 1000 and *γ* = 2 unless otherwise specified.

**Algorithm 3.**
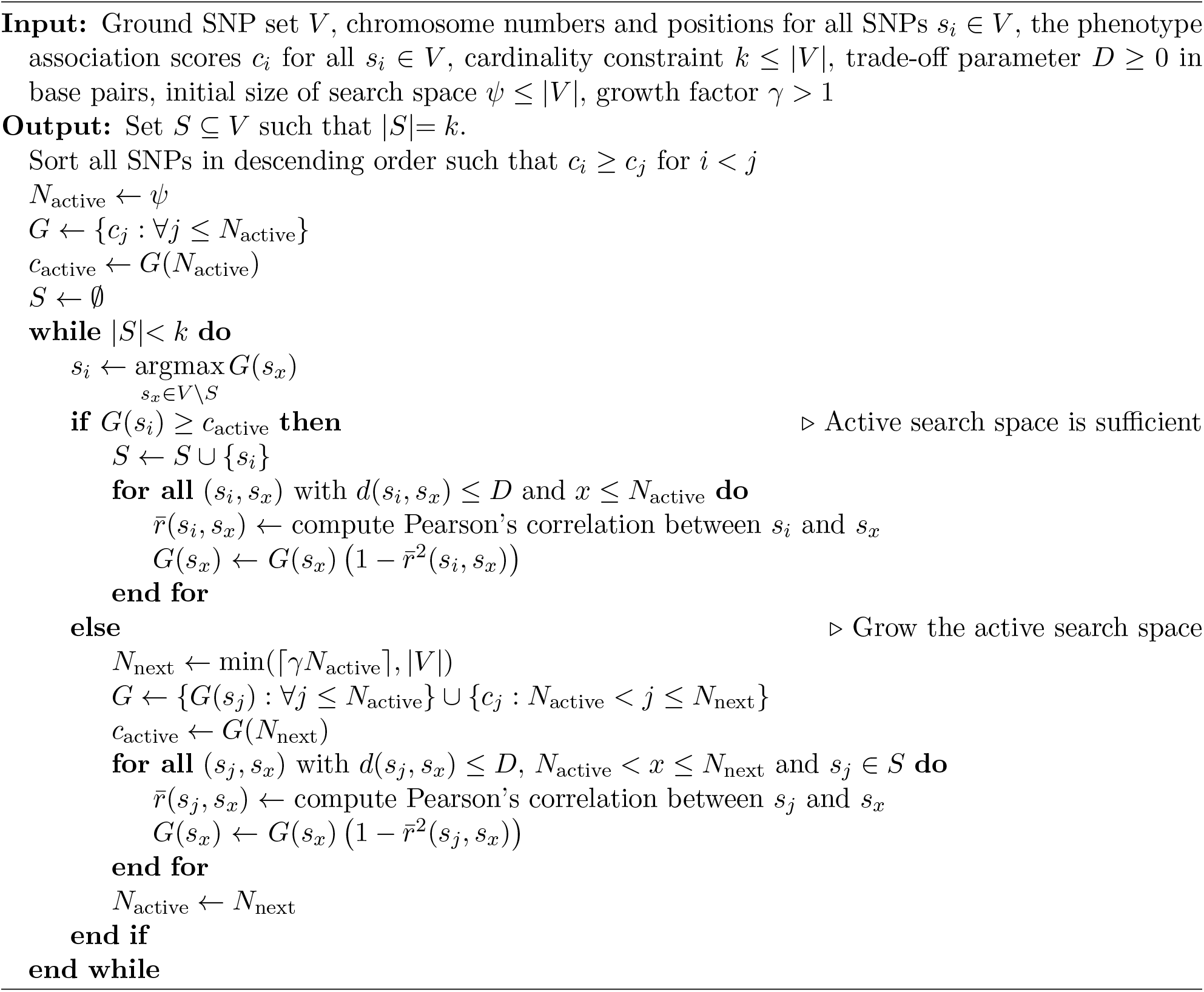
Optimized Macarons Algorithm

## 3 Results

### 3.1 Experimental Setup

#### 3.1.1 Summary of the experiments and the results

First, we investigate the effect of limiting the search of Macarons using intra-chromosomal distance (D parameter) in terms of redundancy and runtime, and whether Macarons can successfully avoid the selection of highly redundant SNPs (Figure 1). Then, we compare Macarons with other SNP selection methods on a small but comprehensive dataset (*AT* dataset with 17 flowering time phenotypes) in terms of their predictivity, runtime and redundancy characteristics (Figures 2–4). Next, we demonstrate that Macarons can seamlessly scale to large datasets with with ~10^7^ variants (in human height dataset). Afterwards, we investigate the utility of avoiding redundancy with Macarons over using a fixed threshold based on individual phenotype association scores on two larger datasets (rice700k and human height) based on 2 different association scores (Figure 5). Finally, we inspect the characteristics of Macarons by visualizing the correlation structure of the selected SNPs while marking the ones near coding regions (Figure 6).

**Figure 1:**
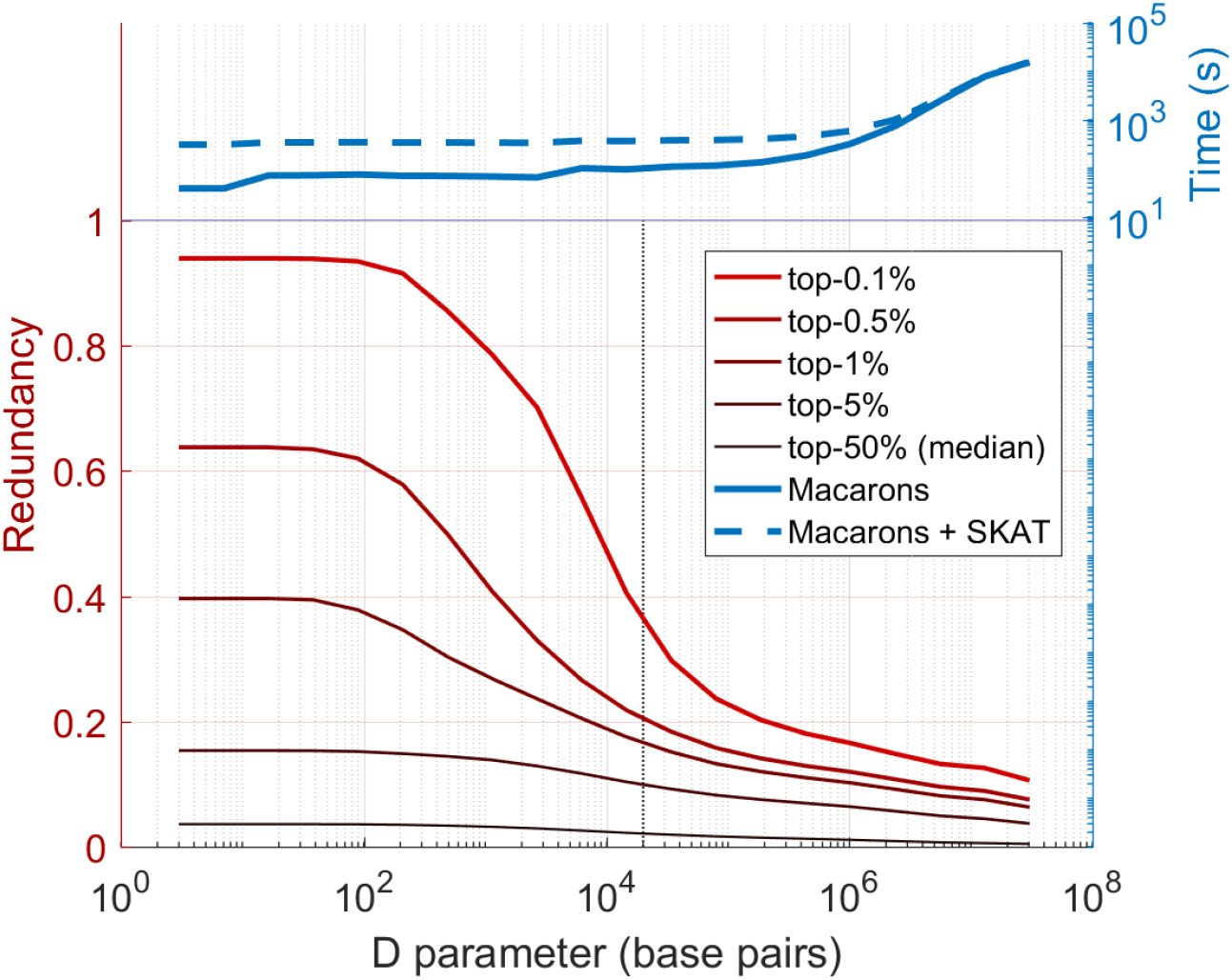
The characteristics of Macarons algorithm with respect to its D parameter in terms of the redundancy between the selected SNPs and the runtime of the algorithm. Redundancy (measured by squared correlation) between all pairs of selected SNPs in AT data for k = 1000. Each line indicates a different percentile for the distribution of redundancy (e.g., top-1% line indicates 1%th most redundant pair). Note that, the lines are averaged across 17 runs corresponding to different flowering time phenotype of AT. The left-most point at the x-axis (for *D* = 1 parameter) corresponds to the baseline method of selecting the highest scoring SNPs (without applying any penalization or regularization). The dotted line indicates *a priori* selected *D* parameter value of 20 kbp. At the top, the blue lines indicate the total runtime (in seconds) to run Macarons for the corresponding *D* parameter (the dashed line additionally includes the time to compute SKAT phenotype association scores).

**Figure 2:**
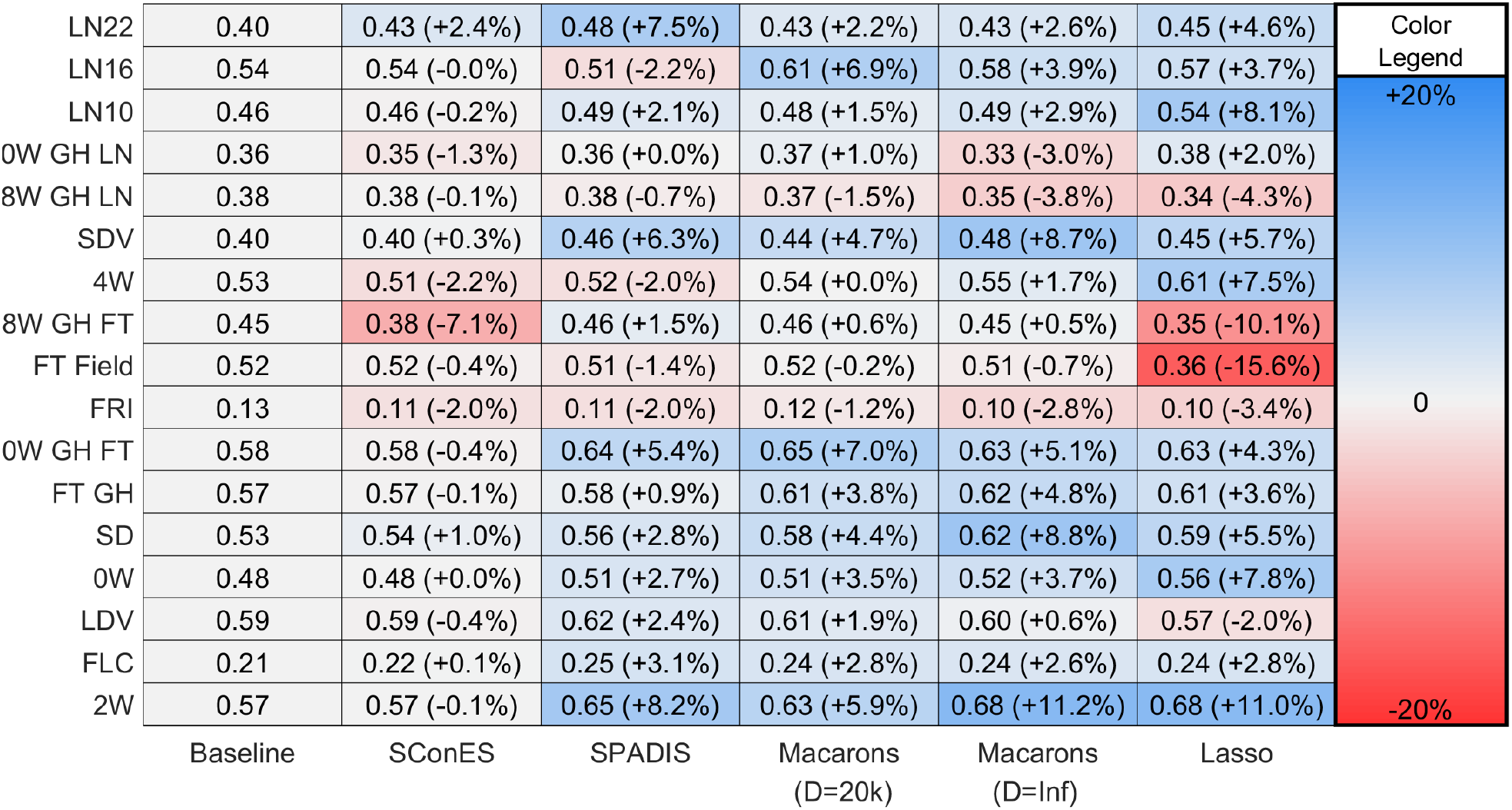
The prediction performances of the SNP selection methods compared to the baseline method for k=1000 selected SNPs. The y-axis corresponds to the 17 flowering time phenotypes and the x-axis corresponds to the SNP selection methods. The leftmost one is a simple baseline method that selects the highest scoring SNPs without any penalization or regularization. Each cell contains two values: (i) The phenotype prediction performance (*R*^2^) for the corresponding method and the phenotype, and (ii) the value in parenthesis shows the difference in *R*^2^ compared to the baseline method as a percentage. Each cell is colored according to their relative improvement compared to baseline: Positive values (indicating improving) are colored blue, and the negative values (indicating decline in *R*^2^ performance) are shown with red colors.

**Figure 3:**
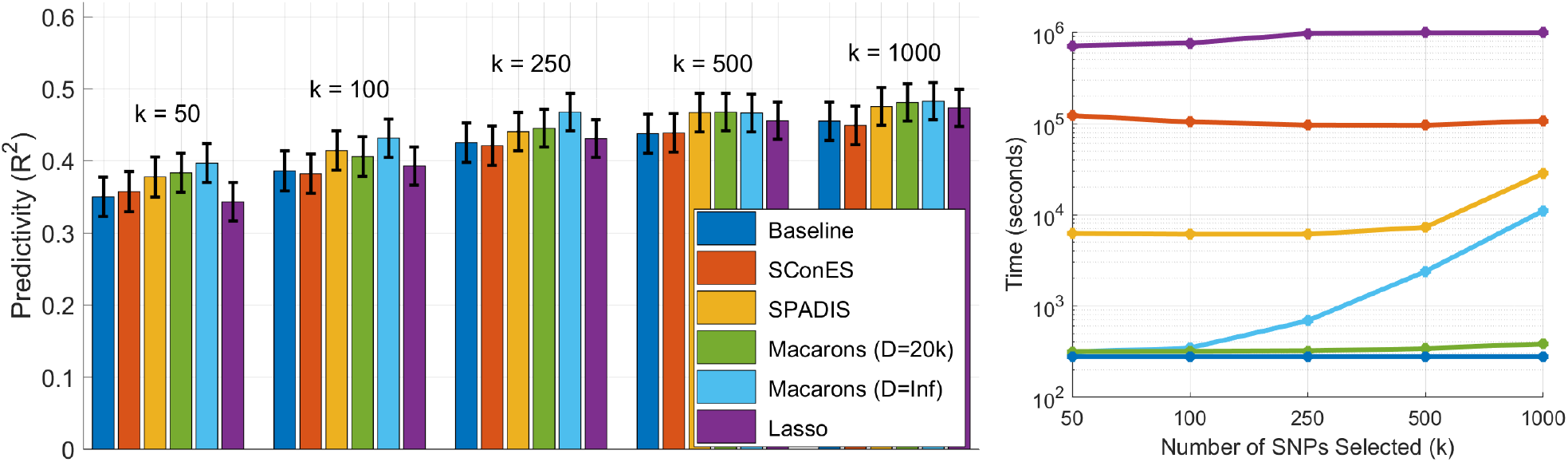
The phenotype prediction performances and runtimes of the SNP selection methods for different number of selected SNPs (indicated by k values). The methods are tested for *k* = 50, 100, 250, and 1000 selected SNPs. (Left) Each colored bar represents a different SNP selection method. The y-axis shows the averaged Predictivity (measured by Pearson’s squared correlation coefficient, *R*^2^) across all 17 flowering time phenotypes. The black lines indicate the 95% confidence interval for the average *R*^2^ performance for the corresponding method and the *k* value. (Right) Each line indicates total time required (in seconds) to run the corresponding method for all 17 flowering time phenotypes.

**Figure 4:**
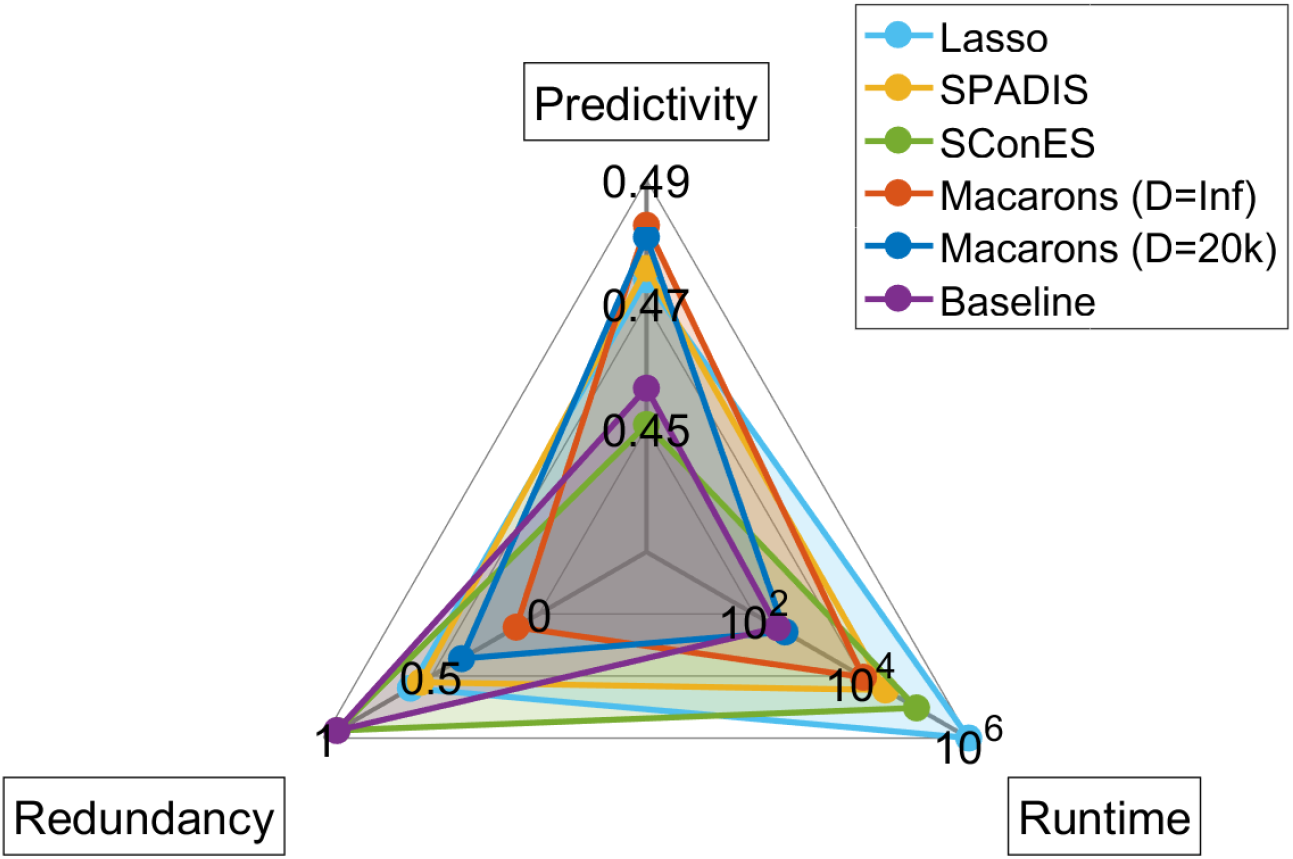
Top-level overview of the characteristics of the SNP selection methods in terms of their predictivity, redundancy, and runtime. The predictivity indicate the average phenotype prediction performance (measured by *R*^2^) of the corresponding method for k=1000 selected SNPs. The redundancy axis indicates the presence of high correlation among the selected SNPs (measured by top-0.1% redundancy: 99.9th percentile of the squared correlation between all pairs of selected SNPs). The time axis (in log-scale) shows the total time required (in seconds) to run the corresponding method for all 17 phenotypes in AT dataset.

**Figure 5:**
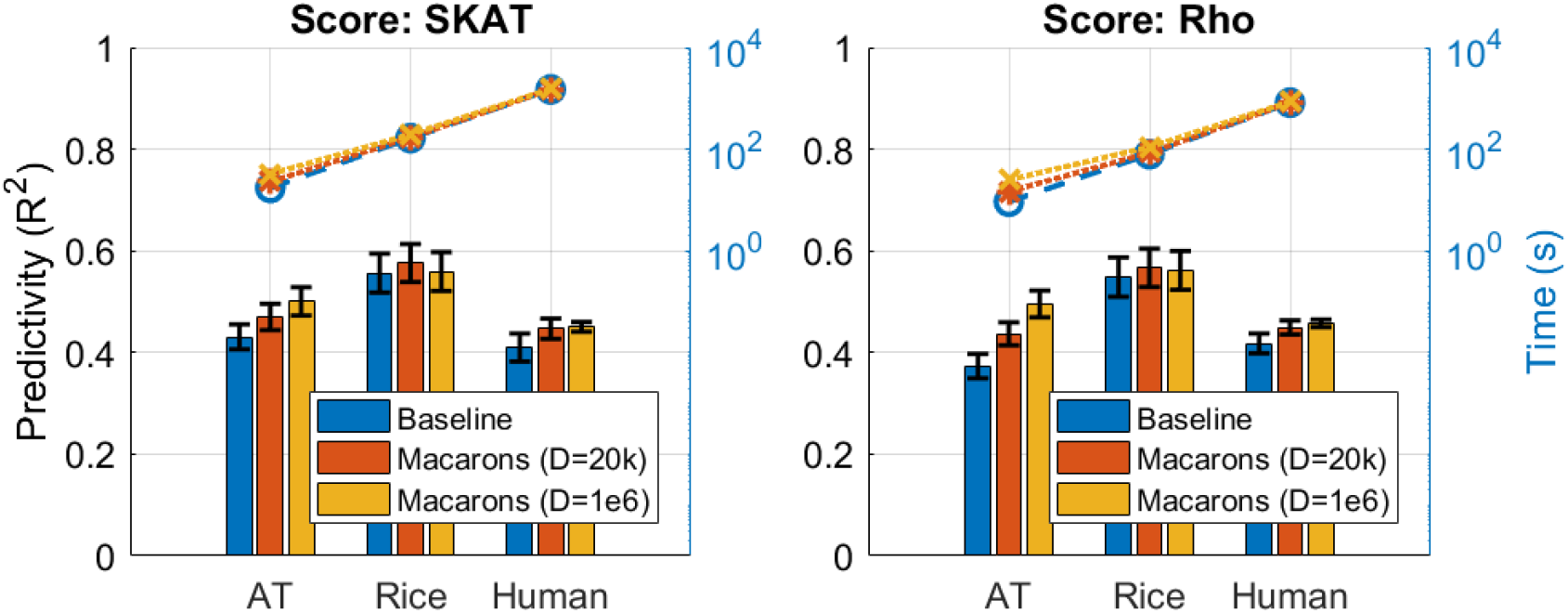
Contribution of Macarons in improving the prediction performance for various datasets and phenotype association scores. The bars represents the predictivity (measured by *R*^2^) of the selected SNPs by the corresponding method for *k* = 1000. The blue bars indicate the prediction performance of baseline method (filtering based on individual phenotype association scores), whereas, red and orange bars indicate the performance of Macarons for various datasets (AT, Rice, Human height) and association scores (SKAT, Rho). The black error bars indicate the 95% confidence intervals. The dashed lines on the top side indicate the runtime of the corresponding method in seconds. Note that, since AT dataset consists of 17 phenotypes, the values shown for predictivity and runtime are averaged across all phenotypes.

**Figure 6:**
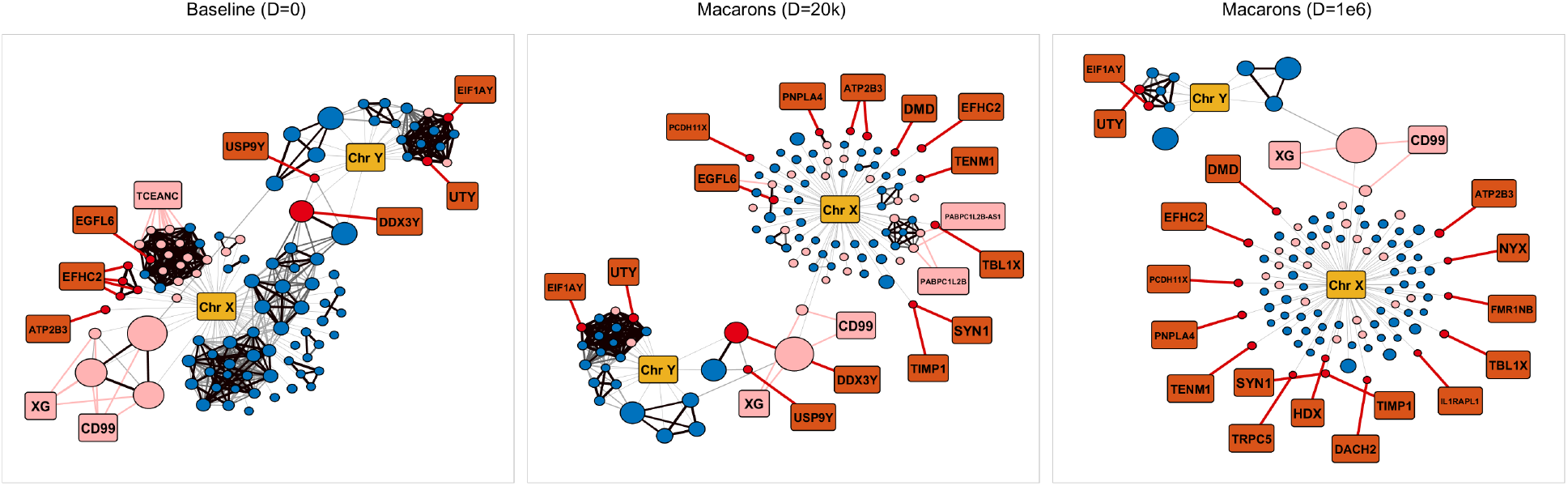
Visualization of the selected SNPs on crowdAI dataset for k=100. Each panel corresponds to a different SNP selection method (Baseline method of selecting top-k SNPs with highest association, Macarons with *D* = 20 kbp, and Macarons with *D* = 10^6^ base pairs). The circles indicate selected SNPs and the rectangles indicate genes (colored red or light red) or chromosomes (colored yellow). Weighted edges between SNPs indicate their redundancy (measured by squared correlation *R*^2^, We include pairs with *R*^2^ ≥ 0.35 and we highlight highly redundant pairs with *R*^2^ ≥ 0.7 with thick lines and black color). The red colored SNPs are within coding region, and light red colored SNPs are within ±20*kbp* around coding region. Similarly, we use red (light red) color for genes with at least one selected SNP in a coding region (around ±20*kbp* of coding region). The sizes of the circles (SNPs) indicate the strength of their individual association with the phenotype (measured by *R*^2^).

#### 3.1.2 Datasets

For a considerable portion of our analysis (e.g., for the comparisons with other SNP selection methods), we use the *Arabidopsis Thaliana* (AT) dataset [2] which provides data for 17 flowering time phenotypes. The availability of multiple phenotype data helps to estimate the variance in phenotype prediction performance more accurately. Also, this is relatively small dataset where the number of samples are between 119 and 180 (depending on the phenotype), and there are 214 051 SNPs before any filtering. Thus, this dataset allows us to test the performance of some methods that would otherwise not scale to larger datasets. In our analysis, we filter out variants with minor allele frequency (MAF) of < 10%, which remains 173 219 SNPs.

As an additional dataset, we use the rice700k data [21] which contains 1145 samples and 700 000 SNPs before filtering. Here, the phenotype is related to the rice grain-length. In our analysis, after applying a MAF < 5% filter, 463 907 SNPs remains. Note that, this is a medium-sized dataset that is roughly 20 times larger than the AT dataset.

As our largest dataset, we consider the human height data^1^ collected from openSNP, which is a crowd-sourced genetic test sharing website [14]. It was prepared by researchers from EPFL as a part of a machine learning challenge on CrowdAI^2^. This dataset contains human height data for 784 individuals and 7 252 636 SNPs. Thus, this dataset is about 10 times larger than the rice700k dataset (and about 200 times larger than AT dataset).

#### 3.1.3 Phenotype association scores

For consistency with the previous results [3, 36], we use Sequence Kernel Association Test (SKAT) [35] to score the individual phenotype association of each SNP, unless otherwise specified (as a part of one of the experiments, we also run our method with another phenotype association measure). While computing the SKAT score, we use the top principal component of the genotype matrix to alleviate the effect of the population stratification [28].

### 3.2 Effect of limiting the search space through intra-chromosomal distance

The premise behind Macarons is to select a complementary set of SNPs while avoiding redundant (correlated) SNPs that are in LD. As we discuss in the methods, the process of taking into account of all redundant SNPs overall requires *k × n* (number of selected SNPs × number of SNPs) correlation estimations from the data, which is both computationally intensive and superfluous since most highly correlated variants tends to be closely located on the genome. To overcome this issue, we limit the search space for correlated SNPs to close intra-chromosomal pairs with maximum distance of *D*, where *D* is an adjustable parameter (unit in base pairs).

Here, we investigate the effect of the parameter *D* on the SNPs selected by Macarons, particularly to examine its effect on the selection of highly correlated SNPs. For this purpose, we select *k* = 1000 SNPs for each of the 17 flowering time phenotypes of AT using Macarons with various *D* parameter values. For each tested value of the *D* parameter, we investigate the distribution of the redundancy for 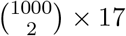 pairs of selected SNPs, in addition to the overall runtime characteristics of Macarons (Figure 1). Note that, since the distribution of the redundancy is greatly skewed (i.e., many pairs with low redundancy, a few with high redundancy), we visualize it using percentile lines (similar to a boxplot) starting from the 50^*th*^ percentile (median) all the way to the 99.9^*th*^ percentile, denoted as top-0.1% redundancy. As it can be seen in Figure 1, with *a priori* selected *D* value of 20 kbps (which is the estimated LD range for AT according to [2]), Macarons can considerably reduce the selection of highly redundant SNP pairs without being bottleneck from a runtime perspective (since the association scores needs to be computed regardless of the *D* parameter or the redundancy calculations). We observe that the redundancy calculations only starts to become a bottleneck after around *D* = 10^6^ base pairs. Next, we investigate whether avoiding the selection of redundant pairs would translate into an improved phenotype prediction performance by comparing the Macarons with other SNP selection methods.

### 3.3 Benchmarking SNP selection methods

#### 3.3.1 Compared Methods

We compare Macarons with the following methods:

- Baseline: A simple greedy approach that selects the top *k* SNPs with the highest individual phenotype association scores. This method becomes equivalent to Macarons when the search space (D) parameter of Macarons is set to 0 (since no redundancy calculations are made and phenotype association scores are not updated in that case). This method considers the association of each SNP independently, thus, serves as a baseline for other SNP selection methods that attempt to take into account of interactions or dependencies between selected SNPs in some manner.
- SConES: A SNP selection algorithm that rewards SNPs according to their individual phenotype association scores of SNPs while employing a connectivity constraint on an SNP-SNP network [3]. It features two parameters λ and *η* that controls the connectivity and sparsity constraints respectively.
- SPADIS, our previous work, rewards SNPs according to their individual phenotype association scores of SNPs while applying a diversity penalty based on the shortest-path distances on an input network [36]. It features three parameters *k* (for number of SNPs selected), *β* (for the strength of penalization) and *D* (for limiting the search range in the network).
- Lasso: A linear regression method with *l*_1_ (lasso) regularization that forces the regression weights of some features (SNPs) to be zero. SNPs with non-zero weights are considered to be selected. It has one parameter λ that determines the strength of regularization and therefore the sparsity (size) of the selected SNP set.

For methods that utilize a SNP-SNP network (i.e., SPADIS and SConES), we use the best performing network Based on the results of previous benchmarkings [3, 36]: Genomic sequence (GS) network (where SNPs that are adjacent on the chromosome are connected) for SPADIS, and Genomic interaction (GI) network (where SNPs that are in the same genomic region as well as the SNPs between interacting genes are connected to form cliques).

Since Macarons has interpretable parameters and does not require a parameter optimization procedure, we tested it for two *a priori* selected *D* values. We choose *D* = 20 kbp as suggested by [2], and we also test *D* = ∞ (which covers the entire chromosome and includes all intra-chromosomal pairs) to see the effect of limiting the search space on phenotype prediction performance.

Note that, to compare phenotype prediction performances of the methods on equal footing, we apply a cardinality constraint *k* on the selected SNP set and compare the results of the algorithms for different values of *k*. To control the number of SNPs selected, the baseline method, SPADIS, and Macarons already has a parameter *k* that we can set directly. On the other hand, SConES and lasso features sparsity parameters that indirectly controls the size of the selected SNP subset. For these methods, we apply a binary search and select the sparsity parameters (*η* for SConES, λ for lasso) that yield the closest number of selected SNPs to the predefined cardinality constraint *k*.

#### 3.3.2 Evaluating phenotype prediction performance

Our testing scheme consists of using a nested cross-validation scheme (outer for evaluation, inner for parameter selection). First, we use 10 cross-validation folds to split the data into training and test samples. For each of the 10 cross-validation folds, we compute phenotype association scores and run the SNP selection methods using training portion of the data, and we predict the phenotype on the test portion using ridge regression. Next, we assess the prediction performance using Pearson’s squared correlation coefficient (*R*^2^) between the predicted and observed (actual) phenotype vectors. Note that, some methods (e.g., SPADIS and SConES) require further cross-validation to tune their parameters. For this purpose, we use a nested-5-fold cross-validation where the training portion of the data is further split into 5 validation folds. On these validation folds, the model’s generalizability to unseen samples is measured by using ridge regression with *R*^2^ and the parameters with highest *R*^2^ are selected. Since Macarons’s parameters are selected *a priori*, it does not require this nested cross-validation procedure.

#### 3.3.3 Comparison of SNP selection methods

Here, to compare the performances of different SNP selection methods, we use the *AT* dataset because it has two main advantages: (i) It contains 17 flowering time phenotypes that allows us to more accurately estimate the phenotype prediction performance (by reporting the averages over all flowering time phenotypes), and (ii) it is relatively small dataset (with ~ 10^2^ samples, ~10^5^ SNPs) which allows us to report results for relatively slow methods (e.g., lasso) that would otherwise not scale to larger datasets. For methods that utilize a phenotype association score (i.e., for all tested methods except lasso), we use SKAT score mainly for consistency with previous benchmarkings that use this dataset [3, 36].

First, we run each method on each of the 17 flowering time phenotypes for *k* = 1000 and assess their 10-fold cross-validated phenotype prediction performance (using ridge regression as the prediction model and measuring by *R*^2^). In Figure 2, we report the prediction performance of the methods relative to the performance of the baseline method (of selecting the top-k SNPs that are most associated to the phenotype individually). Here, we make the following observations:

- SConES does not seem to perform better than the baseline method for predicting the phenotype. We argue that this may be because the network connectivity constraint in SConES reinforces the selection of highly correlated SNPs that are in linkage disequilibrium (LD), which likely pose difficulties for the regression step.
- SPADIS and Macarons (with *D* = 20 kbp) seems to perform quite similarly while both having a higher phenotype performance than the baseline method on most phenotypes.
- Lasso and Macarons (with *D* = ∞, measuring the redundancy of all intra-chromosomal SNP pairs) seems to perform similarly while lasso performs considerably worse than the baseline method on two of the phenotypes.
- For Macarons, using *D* = ∞ to expand the search space over using *D* = 20 kbp does not seem to provide a considerable benefit in phenotype prediction performance for most phenotypes.

Next, in Figure 3, we consider the averaged phenotype prediction performances (*R*^2^, denoted predictivity for brevity) across all 17 phenotypes for various number of selected SNPs (*k* = 50, 100, 250, 500 and 1000). Here, our first observation is that the overall performances of all methods consistently increase as the number of selected SNPs (*k*) is increased. We argue that this is because ridge regression can provide an adequate amount of regularization and improve the predictivity even for relatively large *k* (where *k* > m, the number of samples). Secondly, we observe that the prediction performance of Macarons (for either of the *D* parameter values) is consistently similar or better than all other methods for all *k* values tested.

Additionally, in Figure 3 (right panel), we compare the methods in terms of the total runtime required to run them on the AT dataset (we perform the time measurement on a 40 core machine with Intel(R) Xeon(R) CPU E5-2650 v3 2.30GHz, parallelized on 17 threads for phenotypes). For each method, we report the total CPU runtime with respect to *k* (the reported times include the method runs for all 17 phenotypes and 10-fold cross-validation used for evaluation, as well as calculation of association scores and (if any) cross-validation for parameter tuning). As it can be seen on Figure 3, Macarons with *D* = 20 kbp is at least two orders of magnitude faster than other methods (i.e., SPADIS, SConES, lasso), and compared to the baseline method of using individual association scores for subset selection, improves the predictivity and the redundancy characteristics (Figure 1) of the selected SNP subsets. We also observe that, even though considering all intra-chromosomal pairs (with *D* = ∞) in Macarons does not provide an additional benefit in predictivity over using *D* = 20 kbp for *k* = 1000, the performance of Macarons *D* = ∞ is typically higher than *D* = 20 kbp for lower *k* values. This indicates that, for target subsets of small size, increasing the depth of the search space through *D* parameter might be a more optimal choice.

In Figure 4, we summarize the differences and potential trade-offs between different SNP selection methods by considering three metrics: (i) Predictivity (measured by *R*^2^) for phenotype prediction, (ii) Runtime (total time required to complete our computational experiments, 17 phenotypes with 10-fold each) in seconds, and (iii) Redundancy (measured by top-0.1% redundancy, in a similar manner to the results in Figure 1) that investigates the presence of highly redundant SNP pairs in the selected SNP subset. Overall, Figure 4 suggests that Macarons (*D* = 20 kbp) can offer a good trade-off between different characteristics, with decent predictivity, fast runtime, and a moderate level of redundancy.

### 3.4 Contribution of using Macarons to take dependencies between variants into account

Here, we investigate the effect of Macarons (and avoiding redundancy between the variants) on the characteristics of the selected subsets. First, we compare Macarons with the baseline method (which does not take dependencies into account) in terms of phenotype prediction performance. For this purpose, we benchmark the methods on three datasets (AT, and two larger datasets: rice700k, and human height) and two phenotype association scores: (i) SKAT as done in previous sections, and as an alternative measure (ii) absolute Pearson correlation which is denoted as Rho (ρ). For this analysis, we predict the phenotype using ridge regression on *k* = 1000 selected SNPs and report the performance using *R*^2^. For Macarons, we consider two *D* parameter values : (i) *D* = 20 kilo base pairs as previously done, and (ii) *D* = 10^6^ bp which is approximately the maximum *D* values before Macarons becomes a bottleneck in terms of runtime (according to our analysis on AT data, Figure 1).

As it can seen from Figure 5, we observe a consistent increase in phenotype predictivity when using Macarons across different datasets and association scores although the magnitude of the increase depends on the datasets (e.g., rice dataset exhibits minor differences while the differences in AT are more prominent). In addition, we observe that different association scores result in similar prediction performances. In Figure 5, we also report the overall runtime of the methods (total 10 runs for 10-cross validation folds). As it can seen, Macarons can scale to large datasets (e.g., human height data with ~10^7^ variants) without compromising from runtime (i.e., the computation of phenotype association scores becomes the bottleneck rather than the subset selection).

Next, to elucidate the effect of taking into account of dependency on the characteristics of the selected subset, we visualize the selected SNPs (for *k* = 100 subsets on the human height dataset using *ρ* for phenotype associations) in the form of a correlation network while marking the variants in the coding regions or near ±20 kbp of the coding regions (Figure 6). Our first observation is that there are some highly correlated clusters of variants in the selections of baseline method (can be considered as Macarons with *D* = 0 bp, which considers the variants to be independent). Whereas, these tightly coupled clusters starts to disappear as a higher portion of the dependencies between the variants are taken into account with higher *D* parameter (to the point that there are only a few pairs that are highly correlated for *D* = 10^6^ bp). Interestingly, we also observe that avoiding the redundancies during the subset selection leads to the selection of more variants in coding regions (and more genes with at least one selected variant in their coding region, Supplementary Table 1). Nevertheless, most of the selected variants are not near coding regions, including some of the highly associated ones (Figure 6). Note that, for the human height dataset, all *k* = 100 selected SNPs (regardless of the method used) turn out to be either in chromosome X or Y. This is likely because gender is a strong predictor of height e.g., there is considerable difference in the mean heights of males and females in this dataset (males: 1.79 meters, and females: 1.65 meters).

## 4 Discussion

In order to select a complementary set of SNPs for the prediction of quantitative phenotypes, we develop Macarons, a fast and interpretable model with a simple idea: the joint selection of highly dependent SNPs would be redundant and would not provide complementary information for the prediction of a phenotype.

Overall, this task is known as feature selection in the machine learning literature, and the idea to take redundancy into account is applied extensively. However, most of the established feature selection methods do not scale (from a runtime standpoint) to the SNP selection problem due to the high dimensionality of the GWAS data (e.g., typically up to ~10^7^ variants). Furthermore, such methods suffer from over-fitting since the number of variants is much larger than the number of samples. To overcome these issues, we make simplifying assumptions (as shown in Equation 6 and Equation 10) and limit the search space to intra-chromosomal pairs in close proximity (controlled by a parameter *D* in base pairs, Equation 12).

Our results demonstrate that, with the assumptions and the optimizations in Algorithm 3, Macarons can seamlessly scale to variant sets as large as ~10^7^ in a matter of minutes. We expect that Macarons (with *D* = 20 kbp, or up to *D* = 10^6^ base pairs) can be of practical use in large GWAS studies since it can take into account of the dependencies between the variants without compromising runtime. Overall, it can offer a reasonable trade-off between phenotype predictivity, runtime, and redundancy of the selected subsets.

The intra-chromosomal distance idea and *D* parameter in Macarons can be efficiently generalized to input dependency networks (where the presence of an edge indicates the decision to measure redundancy for that SNP pair, for example, the *D* parameter can be represented as connecting close SNPs as cliques in the network) to limit the search space of the algorithm. We provide a version of Macarons with input dependency network in our implementation though we leave experimentation with it as future work. We expect that this would be useful to take into account of the dependency between variants through more sophisticated models, for example, by considering the 3D structure of the chromosome through Hi-C data.

Macarons can be used in combination with any metric for individual phenotype association (including for dichotomous phenotypes). We expect that Macarons can be especially useful as a part of a multi-stage analysis for performing the initial filtering to reduce the search space, followed by epistasis tests or other subsequent analyses. Overall, the framework we present can be generalized to various other feature selection problems involving high dimensionality within and beyond biomedical applications.

## Code Availability

Source code in Matlab is available at: https://github.com/serhan-yilmaz/macarons

## Footnotes

1 https://zenodo.org/record/1442755

2 https://www.crowdai.org/challenges/opensnp-height-prediction

