## Supplementary Materials for "Uncovering complementary sets of variants for predicting quantitative phenotypes"

Mehmet Koyutürk, A. Ercüment Çiçek, and Oznur Tastan

### 1 Supplementary Text 1

#### 1.1 Properties of the penalty function used in Macarons

Similar to the squared multiple correlation $R^2(S, s_x)$, the simplified penalty function $\bar{R}^2(S, s_x)$ is always between 0 and 1, follows the diminishing returns principle, and has the following properties:

For a given SNP set $S$ and a SNP $s_x \notin S$, the penalization function $\bar{R}^2(S, s_x)$ obeys:

- If $S$ does not include any SNPs correlated with $s_x$, then the penalty for $s_x$ is zero: If $r^2(s_i, s_x) = 0 \ \forall i \in S$, then $\bar{R}^2(S, s_x) = 0$.
- If $S$ includes a single SNP $s_i$, the penalty of $s_x$ is equal to their squared correlation: if $S = \{s_i\}$, then $\bar{R}^2(S, s_x) = r^2(s_i, s_x)$.
- If $S$ includes a SNP $s_i$ perfectly correlated with $s_x$, the penalty for $s_x$ is maximum: if $s_i \in S$ and $r^2(s_i, s_x) = 1$, then $\bar{R}^2(S, s_x) = 1$ (thus, gain function $G'(S, s_x) = 0$).
- The penalty for $s_x$ is monotonically non-decreasing and can only increase when the set grows: $\bar{R}^2(S, s_x) \leq \bar{R}^2(S \cup \{s_i\}, s_x) \ \forall s_i \notin S \cup \{s_x\}$.
- Selecting a SNP $s_i$ uncorrelated with $s_x$ does not affect the penalization of $s_x$: If $r(s_i, s_x) = 0$, then $\bar{R}^2(S \cup \{s_i\}, s_x) = \bar{R}^2(S, s_x)$.
- If $S$ includes two SNPs $s_i$ and $s_j$, the penalization of $s_x$ is equal to the sum of individual squared correlations with $s_i$ and $s_j$ minus their expected overlap: if $S = \{s_i, s_j\}$, then $\bar{R}^2(S, x) = r^2(s_j, s_x) + r^2(s_i, s_x) - r^2(s_j, s_x)r^2(s_i, s_x)$.
- If a new SNP $s_i$ is added to a SNP set $S$, the joint penalization of $s_x$ is increased by the squared correlation of $s_i$ with $s_x$ minus their expected overlap with $S$: If $S' = S \cup \{s_i\}$, then $\bar{R}^2(S', s_x) = \bar{R}^2(S, s_x) + r^2(s_i, s_x) - \bar{R}^2(S, s_x)r^2(s_i, s_x)$.

| | k=100 selected SNPs | | | k=1000 selected SNPs | | |
|---|---|---|---|---|---|---|
| | Baseline (D=0) | Macarons (D=2e4) | Macarons (D=1e6) | Baseline (D=0) | Macarons (D=2e4) | Macarons (D=1e6) |
| Number of SNPs in coding regions | 10 | 14 | 16 | 34 | 113 | 171 |
| Number of SNPs near ±20kbp of coding regions | 31 | 38 | 38 | 96 | 375 | 374 |
| Number of genes with at least one selected SNP | 7 | 14 | 17 | 148 | 197 | 210 |

Table 1: The number of chosen SNPs that lie in coding regions and the corresponding number of unique genes hit for the methods on the human height dataset for k = 100 and k = 1000 selected SNPs by baseline method (corresponding to $D = 0$ parameter of Macarons), Macarons ($D = 20$kbp), and Macarons ($D = 10^6$) when absolute Pearson correlation ($\rho$) is used as the phenotype association score of the SNPs.
